# Time course analysis of the brain transcriptome during transitions between brood care and reproduction in the clonal raider ant

**DOI:** 10.1101/223255

**Authors:** Romain Libbrecht, Peter R. Oxley, Daniel J. C. Kronauer

**Author notes:** co-first authors Correspondence.

## Abstract

Division of labor between reproductive queens and non-reproductive workers that perform brood care is the hallmark of insect societies. However, the molecular basis of this fundamental dichotomy remains poorly understood, in part because the caste of an individual cannot typically be experimentally manipulated at the adult stage. Here we take advantage of the unique biology of the clonal raider ant, *Ooceraea biroi*, where reproduction and brood care behavior can be experimentally manipulated in adults. To study the molecular regulation of reproduction and brood care, we induced transitions between both states, and monitored brain gene expression at multiple time points. We found that introducing larvae that inhibit reproduction and induce brood care behavior caused much faster changes in adult gene expression than removing larvae. The delayed response to the removal of the larval signal prevents untimely activation of reproduction in *O. biroi* colonies. This resistance to change when removing a signal also prevents premature modifications in many other biological processes. Furthermore, we found that the general patterns of gene expression differ depending on whether ants transition from reproduction to brood care or *vice versa*, indicating that gene expression changes between phases are cyclic rather than pendular. Our analyses also identify genes with large and early expression changes in one or both transitions. These genes likely play upstream roles in regulating reproduction and behavior, and thus constitute strong candidates for future molecular studies of the evolution and regulation of reproductive division of labor in insect societies.

## Introduction

The evolution of social life from solitary organisms, one of the major transitions in evolution (1), is best exemplified by eusocial hymenopterans (ants, some bees, and some wasps). At the core of hymenopteran societies lies reproductive division of labor, whereby one or several queens monopolize reproduction while workers perform all the non-reproductive tasks necessary to maintain the colony (2). To better understand the evolution of sociality requires investigating the mechanisms that plastically regulate reproductive and non-reproductive tasks in social insects.

Studies of reproductive division of labor have primarily focused on comparing the queen and worker castes, both at the adult stage and during larval development when caste differentiation occurs (3–6). Such studies have provided valuable insights into the mechanisms regulating the alternative developmental trajectories of queens and workers, and contributed greatly to the elaboration of theories regarding the evolution of sociality (7–12).

However, there are three major limitations associated with the comparison of morphologically distinct queens and workers. First, at the adult stage the two castes not only differ in reproductive status and behavior, but also in morphology, baseline physiology, immunity and lifespan (2, 13, 14). Thus it is difficult to disentangle differences between queens and workers that are actually associated with plastic variation in reproduction and behavior from those associated with other traits. Second, the caste is fixed when females reach adulthood, and thus cannot be experimentally manipulated in adults, making it challenging to establish causality between molecular and phenotypic differences. Third, morphologically distinct queen and worker castes represent the derived state: comparing them does not necessarily provide accurate information on the mechanisms under selection during the evolution of sociality from a totipotent ancestor.

Eusocial hymenopterans are derived from subsocial wasp-like ancestors that alternated between reproductive and brood care phases (8, 15–17). The evolution of sociality involved a decoupling of these phases in different individuals, the queens and the workers, respectively. To understand the evolution of such decoupling requires investigating the molecular mechanisms regulating the transitions between phases. Unfortunately, extant wasp species with a subsocial cycle and progressive provisioning of their larvae are rare tropical species (e.g., *Synagris* wasps in sub-Saharan Africa (18) or *Stenogaster* wasps in southeast Asia (19)) that have not been studied from a molecular perspective because they cannot be experimentally manipulated under controlled laboratory conditions.

The clonal raider ant *Ooceraea biroi* (formerly *Cerapachys biroi* (20)) is a promising model system to study the evolution of sociality because it recapitulates the alternation between reproductive and brood care phases of the subsocial ancestors of eusocial hymenopterans (21, 22). This species has no queen caste, and colonies consist of morphologically uniform and genetically identical workers. Colonies alternate between reproductive phases of ca. 18 days during which workers reproduce asexually in synchrony, and brood care phases of ca. 16 days during which workers have regressed ovaries, forage and nurse larvae (21, 23). Social cues derived from the larvae regulate the transitions between phases: when larvae hatch towards the end of the reproductive phase, they soon suppress ovarian activity and induce brood care behavior in the adults, and when larvae pupate towards the end of the brood care phase the adults begin to activate their ovaries and foraging activity ceases (24, 25). This allows precise experimental manipulation of the cycle by adding or removing larvae of a particular developmental stage at standardized time points during the cycle (Figure 1). At the same time, *O. biroi* affords maximal control over the genetic composition and age structure of experimental colonies, arguably the two most important factors that affect division of labor in social insects (21, 25–27). This study takes advantage of the unique biology of *O. biroi* to investigate the molecular underpinnings of reproductive and brood care behavior, compare general patterns of gene expression between transitions, and identify candidate genes potentially involved in the evolutionary transition from subsocial to eusocial living.

**Figure 1.**
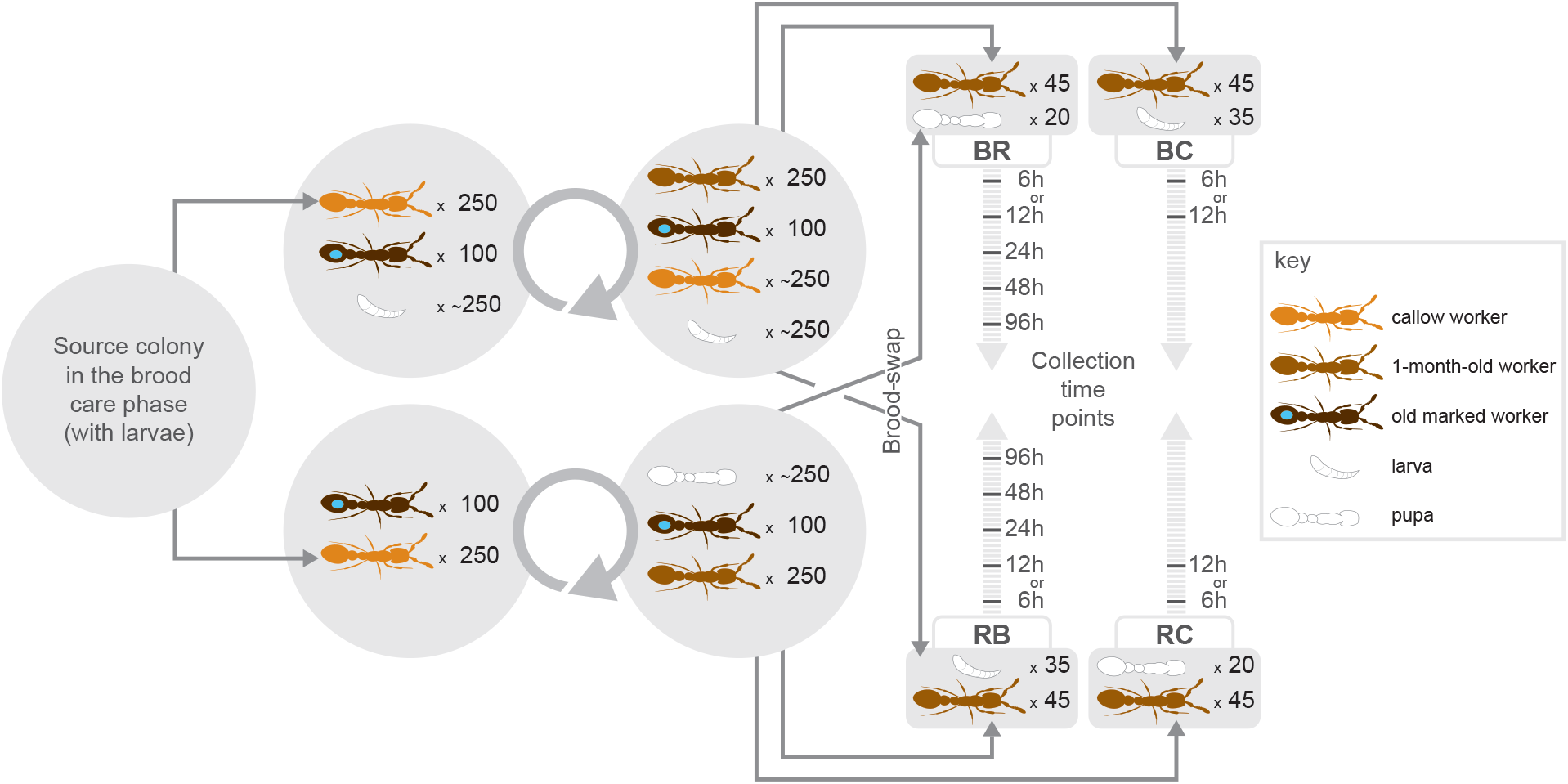
Design of the brood-swap experiment. For each biological replicate, a large source colony in the brood care phase was used to establish two colonies of 250 1-month-old workers and 100 marked ≥ 3-month-old workers. One of these colonies received approximately 250 larvae. After a full colony cycle, each colony contained a complete cohort of brood and workers, and was in either peak brood care phase (with larvae) or early reproductive phase (with eggs and pupae). On the day the first eggs were laid, the 1-month-old workers were subdivided in colonies of 45 workers each. One colony from each phase served as the control colony and was given brood from the mother colony. The remaining colonies received brood from the mother colony in the opposite phase of the cycle, triggering the transition toward the alternative phase. Colonies were subsequently collected 6, 12, 24, 48 or 96 hours post treatment. BR: workers transitioning from the brood care phase to the reproductive phase (after larvae were removed and pupae added); RB: workers transitioning from the reproductive phase to the brood care phase (after pupae and eggs were removed and larvae added); BC: workers from the brood care phase with larvae (brood care phase control); RC: workers from the reproductive phase with pupae (reproductive phase control).

## Results

We experimentally manipulated the presence of larvae in *O. biroi* colonies of age-matched, genetically identical individuals to induce transitions from the reproductive to the brood care phase (hereafter “RB transition”) or from the brood care to the reproductive phase (hereafter “BR transition”). We then collected brain gene expression data from individuals sampled across 5 consecutive time points at 6, 12, 24, 48 and 96 hours post manipulation to evaluate gene expression changes over time in response to changes in brood stimuli (Figure 1). After checking for outliers, we judged the 6-hour time points to mostly reflect a response to recent experimental disturbance, and thus removed them from further analysis (Methods; Supplementary Figure 1).

### Brain gene expression changes when ants transition between phases

We conducted two independent differential expression analyses (one for each transition) that revealed 2,043 genes differentially expressed over time in the RB transition (hereafter “RB-DEGs”) and 626 genes differentially expressed over time in the BR transition (hereafter “BR-DEGs”) (adjusted p-values < 0.05; Methods). These analyses also detected genes with similar patterns of expression in both transitions, which likely stem from experimental manipulations. Thus we conducted a more conservative analysis aiming at identifying genes that showed transition-specific expression changes over time (Methods). We detected 596 genes that showed different expression patterns over time between RB and BR transitions (hereafter “DEGs”; adjusted p-values < 0.05; Supplementary Table 1).

PCA clustering of samples according to brain gene expression segregated samples primarily according to ovary score, and secondarily according to the direction of the transition (Figure 2). Samples that were early in the transition were most similar to their corresponding control samples. Samples that were late in the transition were most similar to the control samples for the opposite transition (i.e., closest to the phase opposite from where they started in the experiment). This shows that our experimental timeline appropriately spanned both transitions from beginning to end, and that brain gene expression is an accurate corollary of the physiological state of *Ooceraea biroi* individuals.

**Figure 2.**
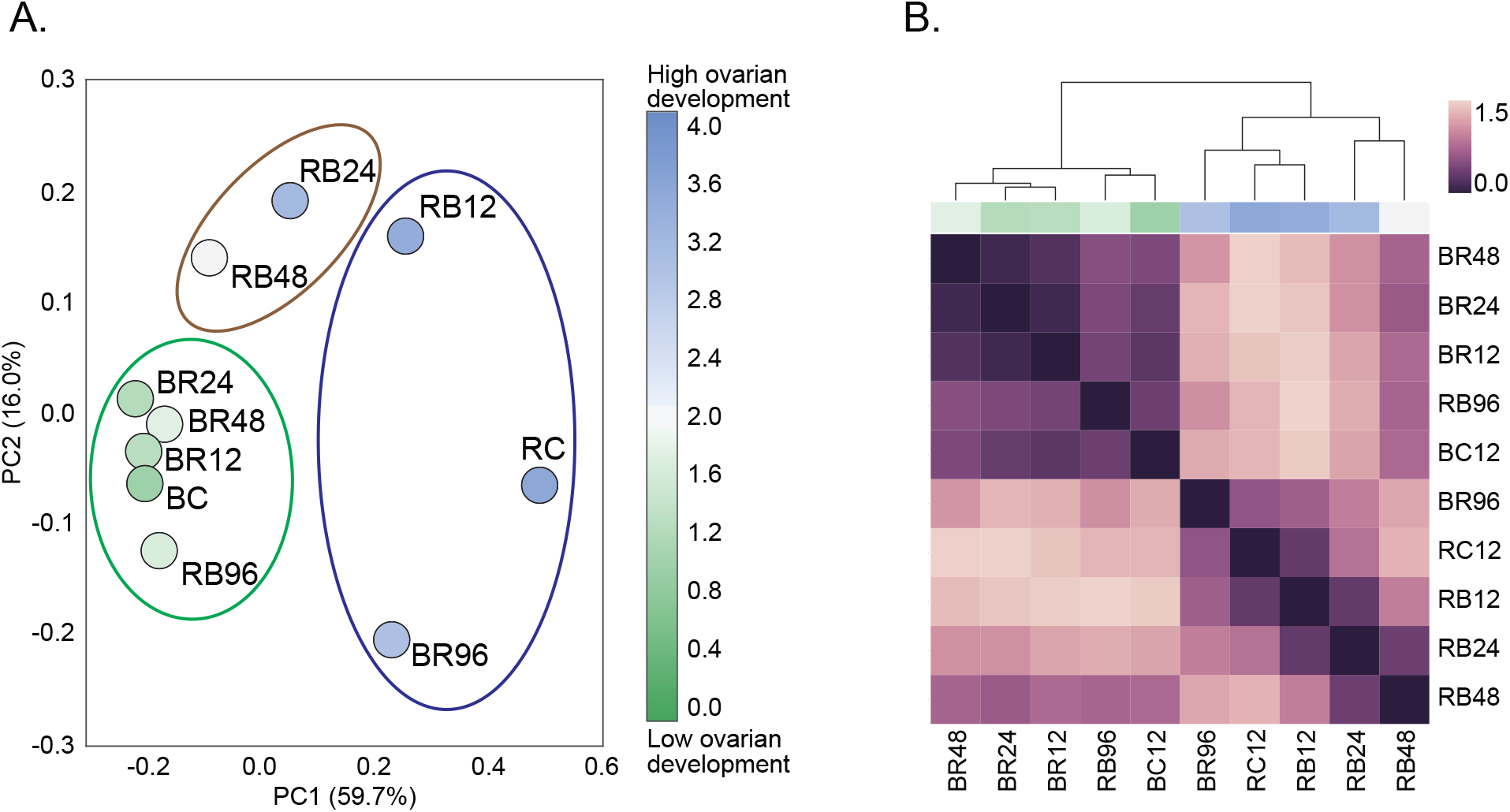
Cluster analysis of samples based on the mean gene expression of each time point, for 596 differentially expressed genes (adjusted p-value ≤ 0.05). A) PCA plot of brood-swap and control samples. Percentages on each axis indicate the proportion of variance explained by the indicated principal component. The blue, brown and green ellipses show the k-means cluster assignment. The color of each sample indicates the average ovary activation score as per Dade, 1994 (70); 0 indicates no signs of ovary activation while 4 indicates fully developed eggs are present. Sample names are as per Figure 1. B) Heatmap showing Euclidean distances between all time points. The dendrogram was constructed using the average distances between time points. The blue and green color bar above the heatmap indicates average ovary activation score, as in A. Sample names are as per Figure 1.

### The timing of gene expression changes differs between transitions

The average distance between samples (Figure 2A and B) indicated a more gradual change in gene expression when transitioning to the brood care phase than when transitioning to the reproductive phase. The unbiased clustering of samples further suggested that changes in gene expression patterns occurred earlier after adding larvae to ants in the reproductive phase than after removing larvae from ants in the brood care phase (Figure 2A). Only samples collected 12 hours after addition of larvae clustered with the control samples for the reproductive phase, while later samples clustered either as an intermediary group (24 and 48 hours), or with the brood care phase controls (96 hours) (Figure 2A). On the other hand, following the removal of larvae, all samples collected before 96 hours clustered with the control for the brood care phase (Figure 2A).

To further test whether gene expression dynamics differed between transitions, we used P-spline smoothing with mixed effects models (28) to fit the gene expression time course profiles into clusters (i.e., groups of co-expressed genes over time). This approach grouped all genes into 76 clusters for the BR transition, and 96 clusters for the RB transition (Supplementary Table 2, Supplementary Figure 2). In order to compare clusters, we also identified their ‘maximum change vector’ (MCV), which is the interval, magnitude and direction of the largest average gene expression change between time intervals. For each transition, we used the MCV values to determine the number of genes showing their maximum change in expression for each time interval. If the timing of gene expression changes was similar in both transitions, we would expect a comparable distribution of such number of genes across time intervals for clusters showing significant changes over time (i.e., clusters enriched for DEGs). Contrary to this expectation, we found that among clusters enriched for DEGs the distribution differed significantly between transitions (χ^2^ = 1217.5, p < 0.00001, Supplementary Figure 3). Consistent with the PCA analysis, most gene expression changes in the BR transition occurred between 48 and 96 hours, whereas changes in the RB transition were weighted towards earlier time intervals (Table 1).

**Table 1.**
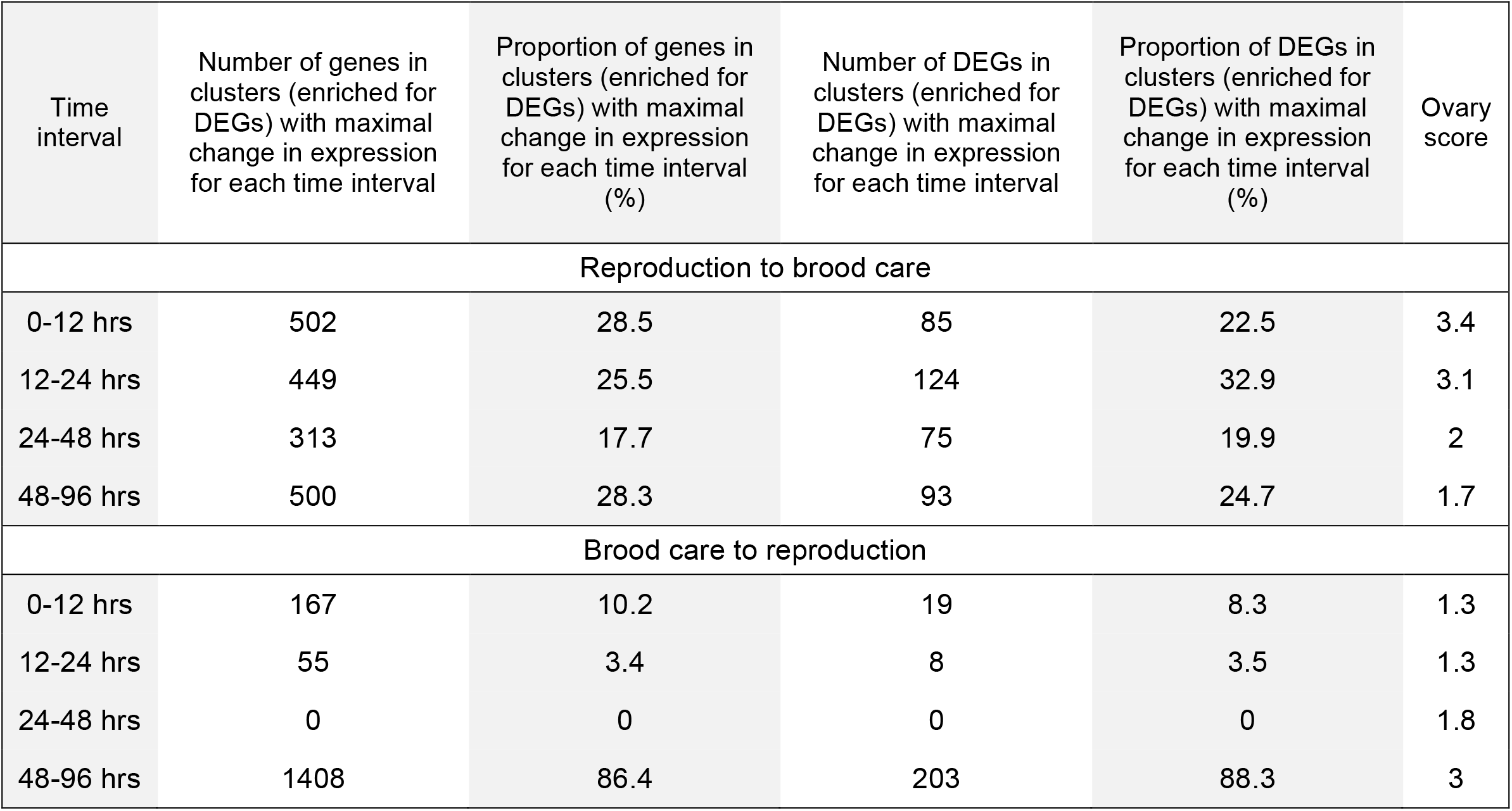
Summary of clusters enriched for DEGs.

| Time interval | Number of genes in clusters (enriched for DEGs) with maximal change in expression for each time interval | Proportion of genes in clusters (enriched for DEGs) with maximal change in expression for each time interval (%) | Number of DEGs in clusters (enriched for DEGs) with maximal change in expression for each time interval | Proportion of DEGs in clusters (enriched for DEGs) with maximal change in expression for each time interval (%) | Ovary score |
| --- | --- | --- | --- | --- | --- |
| Reproduction to brood care |  |  |  |  |  |
| 0-12 hrs | 502 | 28.5 | 85 | 22.5 | 3.4 |
| 12-24 hrs | 449 | 25.5 | 124 | 32.9 | 3.1 |
| 24-48 hrs | 313 | 17.7 | 75 | 19.9 | 2 |
| 48-96 hrs | 500 | 28.3 | 93 | 24.7 | 1.7 |
| Brood care to reproduction |  |  |  |  |  |
| 0-12 hrs | 167 | 10.2 | 19 | 8.3 | 1.3 |
| 12-24 hrs | 55 | 3.4 | 8 | 3.5 | 1.3 |
| 24-48 hrs | 0 | 0 | 0 | 0 | 1.8 |
| 48-96 hrs | 1408 | 86.4 | 203 | 88.3 | 3 |

### The nature of gene expression changes differs between transitions

The independent analyses of the RB and BR transitions revealed a weak overlap between the lists of RB-DEGs and BR-DEGs: only 7.4% (185/2484) of the genes differentially expressed over time in one transition were differentially expressed over time in both transitions. In addition, 55.7% (103/185) of the overlapping DEGs had the same MCV in both transitions, suggesting that their expression changes were a result of experimental manipulation. This suggests that the genes and/or pathways associated with transitioning between phases are specific to each transition.

The gene co-expression clusters further corroborate this finding. Constructing a network from cluster membership in both transitions revealed a highly connected, homogenous network (Supplementary Table 3), showing that most genes were co-expressed with different genes in each transition. This is similarly illustrated by cluster enrichment for Gene Ontology (GO) terms. We found 27 enriched clusters (including four clusters enriched for DEGs) for the BR transition, and 35 (including seven clusters enriched for DEGs) for the RB transition (Supplementary Table 2). Among clusters enriched for DEGs, only 6.9% (2/29) of the GO terms associated with one transition were also associated with the other transition (Supplementary Figure 4).

Furthermore, the expression patterns of genes that were co-expressed with the same genes in both transitions were inconsistent with a symmetrical molecular regulation. We identified all conserved co-expression clusters in the network (i.e., clusters whose members were more similar between transitions than expected by chance) (Methods, Supplementary Table 3). If the primary molecular mechanisms regulating phase transitions were reversible, then co-expressed genes would show expression changes in opposite directions in each transition. In that case, we would expect network edges that link clusters of genes regulated in opposite directions between transitions to represent a higher proportion of edges in the conserved network (with only nonrandom connections) compared to the complete network (which includes random connections). Contrary to this expectation, edges linking clusters of genes showing opposite patterns of expression changes over time between transitions were less frequent in the conserved network (24.4%) than in the complete network (45.8%; χ^2^ = 22.4, p < 0.00001; Supplementary Figure 3).

### Using the time course data to identify candidate genes

Ranking the 596 DEGs according to their change in expression between the control and the 96-hour time point for each transition allowed us to identify genes most likely to be involved in the molecular regulation of one or both transitions (Supplementary Table 4). This includes genes that encode proteins with neuroendocrine functions (*Vgq* or *queen vitellogenin*), neuropeptides (*insulin-like peptide 2, neuroparsin-A*) and neuropeptide receptors (*leucine-rich repeat-containing G-protein coupled receptor 4*), enzymes involved in neuropeptide processing (*carboxypeptidase M, aminopeptidase N*), neurotransmitter receptors (*glycine receptor subunit alpha 2*), proteins involved in neurotransmission (*synaptic vesicle glycoprotein 2C*, three *kinesin-like proteins*), neuronal function (*leucine-rich repeat neuronal protein 2, trypsin inhibitor, gliomedin*), transcription (*hunchback, transcription termination factor 2, speckle-type POZ protein B, zinc finger BED domain-containing protein 1, lymphoid specific helicase*), and protein synthesis and modification (*peptidyl-prolyl cis-trans isomerase D, hyaluronan mediated motility receptor, alpha-(1,3)-fucosyltransferase 6*). The expression patterns for some of these candidate genes are shown in Figure 3.

**Figure 3.**
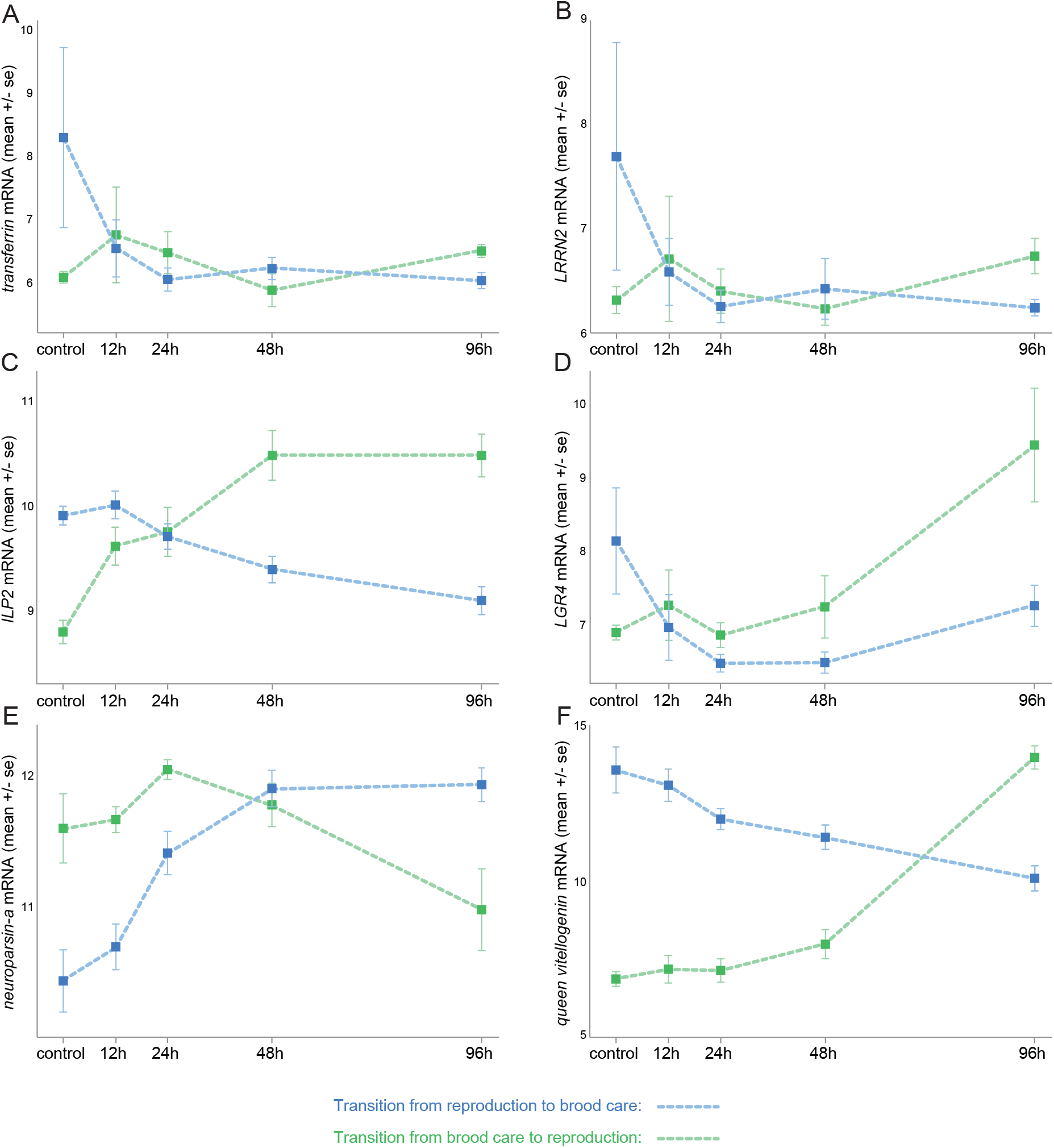
Select candidate genes for the regulation of the transitions between brood care and reproduction. The plots show the expression changes over time after adding larvae (RB transition; blue) or removing larvae (BR transition; green) from the colonies. A) *transferrin*; B) *LRRN2 (leucine-rich repeat neuronal protein 2*); C) *ILP2* (*insulin-like peptide 2*); D) *LGR4* (*leucine-rich repeat-containing G-protein coupled receptor 4*); E) *neuroparsin-a; F*) *queen vitellogenin*. Gene expression is shown as variance-stabilized transformed read counts (which approximate log2-transformed read counts).

In addition, we identified genes for which expression changed early in the transitions, i.e. genes that likely function upstream in the molecular processes regulating the transitions. For each transition we generated a list of genes with highest change in expression between the control and the 12-hour time point (Supplementary Table 4). These lists include candidate genes with early changes in the RB transition (*hunchback, alpha-(1,3)-fucosyltransferase 6*), in the BR transition (*insulin-like peptide 2, glycine receptor subunit alpha 2, transcription termination factor 2, hyaluronan mediated motility receptor, annulin*), or in both transitions (*leucine-rich repeat-containing G-protein coupled receptor 4, leucine-rich repeat neuronal protein 2, transferrin*). Most of the genes for which expression changed early in the transition were also identified due to their strong change in expression between the control and the 96-hour time point (Supplementary Table 4).

### Both transitions are regulated by overlapping sets of transcription factors

For each transition, we tested whether gene clusters were enriched for transcription factor binding sites (TFBSs). We used the JASPAR database to identify 27 clusters (including four clusters enriched for DEGs) in the BR transition and 12 clusters (including four clusters enriched for DEGs) in the RB transition that were significantly enriched for TFBSs (Supplementary Table 2). A number of transcription factors were repeatedly associated with clusters enriched for DEGs, and were present in both transitions (Supplementary Table 5). Of particular note, in each transition there was only one cluster enriched for a single TFBS, and in both cases it was for the *forkhead* binding site. We identified all genes with at least one highly conserved binding site for *forkhead* (Methods) to show that these genes cluster samples according to ovary activation and chronological distance (Supplementary Figure 5), which is consistent with *forkhead* being involved in the regulation of both transitions.

## Discussion

Colonies of *O. biroi* alternate between brood care and reproductive phases, and our time course analyses of the brain transcriptome reveal that the transitions from brood care to reproduction and from reproduction to brood care involve distinct overall patterns of gene expression changes. The timing of brain gene expression changes after manipulating social cues differs between transitions. The addition of larvae leads to a rapid change in gene expression, whereas larval removal results in a much slower change. Inappropriately-timed production of eggs incurs individual and colony-level fitness costs. At the individual level, eggs laid in the presence of larvae are eaten, wasting the resources taken to produce them. Furthermore, individuals with active ovaries are aggressed and eventually killed by nestmates (25). Such policing behavior has been hypothesized to minimize colony-level costs because unsynchronized egg-laying would jeopardize the colony cycle (29). Such fitness costs will exert selective pressure on the regulation of reproductive physiology (30): in line with our findings regulatory mechanisms should be slow to activate ovaries, and quick to suppress or reverse egg production.

In addition, our results are consistent with larval cues acting as a reinforcement signal for brood care and for inhibition of reproduction, because the removal of the brood signal is accompanied by a delay in gene expression and physiological adjustments. Such a delay is necessary in *O. biroi* to prevent premature transitioning to reproduction, such as during foraging, when some individuals frequently exit the nest during the brood care phase and are thus only sporadically exposed to larval cues. Comparable resistance to change has been observed in other species and contexts. In behavioral sciences, the resistance to change in behavior after removal of a stimulus has been compared to the inertial mass (31), and applied to behaviors as diverse as drug addiction in humans (31) or food-reinforced behaviors in birds (32). Physiological regulations are also subject to resistance to change. For example, physiological changes that occur in rats in response to a stressful stimulus (e.g., cold temperature or low oxygen) take several days to return to baseline levels after stimulus removal (33, 34). This pattern of rapid response to stimulus exposure but slow response to stimulus removal also parallels the adaptation and deadaptation rates seen in many molecular systems, such as the cAMP-mediated cGMP response inducing cell aggregation in the slime mold *Dictyostelium discoideum* (35).

We expected that the *O. biroi* colony cycle would be regulated by discrete gene networks in which expression is coordinately and symmetrically up- or down-regulated during transitions between phases. However, neither the differential expression nor the network analyses found substantial overlap in gene membership between transitions. In other words, the sequence of gene expression changes that is associated with the transition to the reproductive phase is not the reverse sequence of gene expression changes associated with the transition to the brood care phase.

Finding different general expression patterns between transitions does not necessarily imply that individual genes or groups of genes cannot play a regulatory role in both transitions. In fact, some of the strongest candidate genes identified here are involved in both the BR and the RB transition (see below). Our differential expression analysis revealed that genes with some of the highest expression changes over time have neuroendocrine, neuronal and gene regulatory functions, and regulate neuropeptide signaling and neurotransmission. Among these genes we have identified some candidate genes for the regulation of reproductive division of labor (Figure 3) by focusing on those that show some of the largest and/or earliest changes in expression after manipulating the presence of larvae.

The gene *transferrin* (Figure 3A) shows large and early changes in expression in both transitions. Moreover, *transferrin* shows caste-specific expression in multiple species of social insects. In the ant *Temnothorax longispinosus* and in the wasp *Polistes canadensis*, whole-body RNA sequencing revealed higher expression in queens compared to workers (36, 37). However, the gene likely has tissue-specific and context-dependent expression patterns and functions (38). While in insects the protein encoded by *transferrin* transports iron into the eggs, reduces oxidative stress, and interacts with the vitellogenin and juvenile hormone pathways (39), its role in the brain remains poorly understood.

The genes *leucine-rich repeat neuronal protein 2* (*LRRN2;* Figure 3B) and *glycine receptor subunit alpha-2-like* (*GLRA2*) show early and opposite expression changes between transitions. They have not been studied in social insects but are involved in neuronal function and neurotransmission in vertebrates. *LRRN2* encodes a protein involved in the development and maintenance of synapses in vertebrate brains (40), and *GLRA2* encodes a subunit of the receptor of the neurotransmitter glycine (41).

Another candidate gene identified in our study is *insulin-like peptide 2* (*ILP2*; Figure 3C), a neuropeptide that belongs to the insulin signaling pathway, which is a conserved pathway that regulates nutrition, fertility and longevity in animals (42, 43). Insulin signaling, together with the juvenile hormone and vitellogenin pathways (44–46), is involved in caste determination and division of labor in social insects (7, 44, 47–50). Interestingly, *ILP2* shows one of the earliest responses to the removal of larvae (in the BR transition), thus suggesting that *ILP2* plays an upstream role in the regulation of reproduction. Another candidate for an upstream regulatory role, in this case in both transitions, is the gene *leucine-rich repeat-containing G-protein coupled receptor 4* (*LGR4*; Figure 3D). It encodes a G-protein coupled receptor predicted to bind relaxin-like peptides (51), which belong to the insulin family, together with insulin-like peptides and insulin-like growth factors (52). Together, these genes suggest an important role for the insulin signaling pathway in linking changes in social cues to reproductive changes.

The expression of the neuropeptide *neuroparsin-a* increases gradually when transitioning to brood care (Figure 3E), which is consistent with neuroparsins having anti-gonadotropic roles (53). To our knowledge, neuroparsins have not been reported to regulate caste differentiation and reproductive division of labor in ants, but they interact with the vitellogenin and insulin signaling pathways (53), and are involved in regulating another well-characterized instance of phenotypic plasticity, the phase transitions in locusts (54).

*Queen vitellogenin* (Figure 3F) is differentially expressed between reproductive and non-reproductive castes in multiple species of ants, bees, wasps and termites (10, 21, 36, 47, 55–60). This gene encodes the yolk protein precursor vitellogenin, which is instrumental to egg formation. In formicoid ants, the *vitellogenin* gene has been duplicated, and some gene copies have been co-opted to regulate non-reproductive functions such as behavior (10). The *queen vitellogenin* gene in *O. biroi* clusters with the gene copies having the ancestral function of regulating reproduction in other species (21). The *worker vitellogenin* gene, which clusters with genes regulating behavior in *Solenopsis invicta* (56) and *Pogonomyrmex barbatus* (10), was not one of the DEGs in this study. Interestingly, the changes in *queen vitellogenin* expression mirror the ovarian development and overall alterations of the transcriptome: *queen vitellogenin* displays a gradual and early decrease during the RB transition but a sharp and delayed increase during the BR transition (Figure 3F).

The protein vitellogenin is typically synthesized in the fat body, secreted into the hemolymph and transported into the developing oocytes (61). However, in some cases caste-specific *vitellogenin* expression has been detected in the head (21, 47) or in the brain (57). In addition, the vitellogenin protein has been localized in the honeybee brain, suggesting that vitellogenin also has a neuroendocrine function (62). Here we show that *vitellogenin* gene expression in the brain is correlated with changes in reproductive physiology, and is consistent with a regulatory function in the brain. Although *queen vitellogenin* shows one of the highest changes in expression in both transitions, such changes occur rather late after manipulating social cues. This suggests that the role of vitellogenin in the brain is likely to be downstream of earlier molecular changes associated with each transition.

Recent studies have proposed that changes in gene regulatory mechanisms were associated with the evolution of eusociality (63, 64). In our study, many DEGs that showed early changes in gene expression have gene regulatory functions such as the onset (*hunchback*) and termination (*transcription termination factor 2*) of transcription, as well as the synthesis (*PPID, annulin*), glycosylation (*alpha-(1,3)-fucosyltransferase 6*) and phosphorylation (*hyaluronan mediated motility receptor*) of proteins. In addition, gene clusters enriched for DEGs were also frequently found to be enriched for genes with certain transcription factor binding sites. This suggests complex transition-specific gene expression and regulation, affected by multiple transcription factors. Nevertheless, genes that are putatively regulated by a few transcription factors exhibit predictable patterns of regulation. For example, the expression of genes associated with *forkhead* transcription factor binding sites provided significant predictive power as to the physiological state of an individual. Interestingly, *forkhead* transcription factors regulate reproduction in other insect species. For example, knocking down *forkhead* transcription factors in the yellow fever mosquito *Aedes aegypti* and the brown planthopper *Nilaparvata lugens* reduced offspring production and the activity of the vitellogenin pathway (65, 66). In addition, *forkhead* plays a role in the regulatory network of salivary glands in insects (67), which include the mandibular glands that produce caste-specific compounds in honeybees (68). Interestingly, the promoter region of *forkhead* shows a depletion of TFBS in ants compared to solitary insects, which may have facilitated *forkhead* pleiotropy and its implication in caste-specific regulatory networks (64). The decoupling of brood care and reproductive phases in different female castes during the evolution of eusociality was associated with the co-option of gene function and regulation (8). Our findings suggest that transcription factors such as *forkhead* may be among the regulatory elements responsible for the co-option of gene regulatory networks during this evolutionary transition.

Studies of caste-specific gene expression patterns in eusocial hymenopterans, combined with cross-species comparisons, have led to the development of multiple hypotheses for the molecular mechanisms associated with the evolution of eusociality (7–12). While our study was not designed to investigate their validity, the asymmetry observed in the gene networks between transitions in *O. biroi* indicates that such hypotheses must be able to account for the different molecular trajectories underlying the transitions between brood care and reproductive phases in the subsocial ancestors of eusocial hymenopterans. This suggests that comparisons between the queen and worker castes and identification of single genes and/or gene networks that are differently expressed between them can only provide a partial understanding of the molecular mechanisms behind the evolution of eusociality.

Assuming that the colony cycle of *O. biroi* indeed represents a partial reversal to the lifecycle of the subsocial ancestor of ants and possibly other eusocial hymenopterans, one parsimonious way to compartmentalize such a cycle would be to disrupt the transition to brood care in response to larval cues in a subset of individuals, which would then act as queens. Given that these queens would now lay eggs continuously, any additional females that emerge at the nest would immediately and permanently be exposed to larval cues and thus locked in the brood care phase of the ancestral cycle. This would then give rise to reproductive division of labor, which could be acted upon by natural selection, driving continued divergence in fertility, and ultimately leading to eusociality. In this study we report that patterns of gene expression changes over time differ between the transition to brood care and the transition to reproduction in *O. biroi*. Our results are therefore not consistent with the transitions being regulated by mirrored sequences of gene expression changes in a pendular manner. On the contrary, patterns of gene expression appear to be circular, with the involvement of transition-specific sets of genes. This implies that, on a molecular level, the transition to brood care could have been disrupted in a variety of ways without affecting the reverse transition. However, especially given our finding that exposure to larval cues entails rapid and large-scale changes in brain gene expression, we would assume that this disruption happened early and upstream in the gene expression cascade. Our time-course data allowed us to identify molecular candidate pathways that respond rapidly to larval cues and are therefore likely upstream of the longer-term behavioral and physiological responses. These constitute prime candidates, both for broad comparative analyses across social hymenopterans, as well as for functional experiments in *O. biroi* and other species.

## Methods

### Biological samples

Source colonies (Figure 1) were derived from two separate clonal lineages: MLL1 and MLL4 (69). Clonal lineage and source colony identity are recorded for all RNA sequencing libraries, which are uploaded to the Short Read Archive (project accession PRJNA273874). Large source colonies in the brood care phase were used to establish two experimental colonies each (250 one-month old workers and 100 ≥ 3-months old workers), one of which received approximately 250 larvae. After a full colony cycle, each colony contained a complete cohort of brood and workers, and was in either peak brood care phase or early reproductive phase. On the day the first eggs were laid in the reproductive phase colony, the one-month old workers were subdivided into colonies of 45 workers. One of these colonies from each phase served as the control colony and was given brood from the colony the workers were derived from (i.e. larvae for the brood care phase control and eggs and pupae for the reproductive phase control). The remaining colonies received brood from the colony at the opposite stage of the cycle (sub-colonies originally in the reproductive phase received larvae and *vice versa)*, thereby inducing the transition toward the opposite phase. All colonies with larvae were fed every 24 hours, immediately after samples for the respective time points had been collected. Colonies were collected 6, 12, 24, 48 or 96 hours after experimental manipulation. This process was repeated eight times: four times without the 6-hour time point (and a control collected after 12 hours) and four times without the 12-hour time point (and a control collected after 6 hours). After looking for outliers, we removed all samples collected after 6 hours (see details below), thus resulting in 4 biological replicates for the controls and 8 biological replicates for all other time points. Source and experimental colonies were kept at 25°C and 60% humidity, and when in the brood care phase were fed frozen *Solenopsis invicta* brood.

### Sample collection and RNA Sequencing

At the specified time for each colony, all ants were flash frozen and subsequently stored at - 80°C. Ovaries and brains were dissected in 1x PBS at 4°C. Ovary activation was measured according to Dade *et al*. (70). Brains of individuals with two ovarioles were transferred immediately to Trizol, and once ten brains from a colony were pooled, the sample was frozen on dry ice.

RNA was extracted with RNEasy column purification, as explained in Oxley *et al*. (21). Clontech SMARTer low input kits were used for library preparation and RNA sequencing was performed on a HiSeq 2000, with 100 bp single end reads. Sequencing batches included all time points for both transitions of any given colony, for two source colonies at a time.

### Identification of outlier samples

967 genes had more than 2-fold change in expression between sample means. These genes could be used to observe the general pattern of sample clustering, before filtering for genes that showed differences between our intended treatment classes (Supplementary Figure 1).

All 6-hour samples (controls and treatments) clustered more closely with each other than with their respective (expected) transition groups. Looking at individual gene expression time courses, it was clear that the 6-hour time points frequently deviated wildly from the other time points. This suggests that the majority of gene expression changes observed in the 6-hour time points was induced by the experimental disturbance. However, removing the 6-hour time points could prevent us from detecting genes that legitimately changed as a result of the actual brood-swap, instead of the experimental manipulation. We therefore looked at the change in sensitivity and specificity of the experiment after removing the 6-hour samples from the analysis.

Removing the 6-hour time points reduced the number of genes with ≥ 2-fold difference by 335. 51% of these 335 genes were differentially expressed between 6- and 12-hour control samples of the same phase, and were therefore *a priori* likely to be false positives. 73 genes were expressed ≥ 2-fold between 6-hour control and treatment samples, and were therefore potentially genes regulated by the change in brood stimuli. Of these 73 genes, only 5 were not present in the 632 genes still identified as having ≥ 2-fold differences after removal of the 6-hour time points (Supplementary Figure 1). If these genes were real target genes, we would only lose 6.8% of the early-responding genes. Removing the 6-hour time points as outliers therefore increased the specificity of our differential expression analysis, with negligible loss of sensitivity.

### Identification of differentially expressed genes

Fastq reads from all samples were aligned to the *Ooceraea biroi* genome (NCBI assembly CerBir1.0) using STAR (default parameters). HTSeq was then used to determine the number of reads aligned to each gene (NCBI *<u>Cerapachys biroi</u>* Annotation Release 100). DESeq2 was used for differential gene expression analysis.

To analyze each transition separately, we contrasted the following models in DESeq2:

> full model: colony + bs(time, df=3)
>
> reduced model: colony

using the bs function from the splines library (v. 3.2.3) in R for evaluating the spline function of all time points (controls coded as time 0). This contrast identified genes with a significant change at any point in time, not just genes significantly different from the control samples. This analysis was run for both BR and RB transitions.

To account for the effects of experimental manipulation, the following models were contrasted:

> full model: colony + transition + bs(time, df=3) + transition: bs(time, df=3)
>
> reduced model: colony + transition + bs(time, df=3)

This model contrast identified the genes that were differentially expressed over time, after accounting for the differences in gene expression between reproduction and brood care phases. Without using the spline function, we could simply be comparing gene expression at each time point to “time 0” (i.e., the control samples). However, this would not reveal genes whose expression changed temporarily, before returning to their baseline value.

We identified only those genes with a significant time by transition interaction. It has been shown that expression of certain genes can have opposing effects, depending on the context (71). Genes that show significant change in expression over time, but no significant interaction with phase, may therefore still be important in regulating transitions between phases. However, such genes are confounded with, and cannot be disentangled from, genes that are expressed as a stress response resulting from the brood-swap experimental procedure, and we therefore decided to ignore them in our present analyses.

### Clustering of gene expression time courses

We clustered the samples using P-spline smoothing and mixed effects models according to the algorithm by Coffey *et al*. (28). To determine the optimal number of clusters for each transition, we calculated the BIC score for all even cluster sizes between 2 and 120 clusters (Supplementary Figure 2). We selected the smallest cluster size of the lower BIC values that did not precede a higher BIC value (Supplementary Figure 2).

### Enrichment analyses for expression clusters

Transcription factor binding site (TFBS) enrichment of each cluster was determined with Pscan, using the available position weight matrices from the JASPAR database. Assessment of clusters for enrichment for DEGs and GO terms was determined using Fisher’s exact test followed by Benjamini & Hochberg (72) false discovery rate adjustments. To identify all *O. biroi* annotated genes with *forkhead* TFBS, we used the R packages TFBSTools and biostrings, with the position weight matrix for *Drosophila* from the JASPAR database, and a 95% minimum score for matching.

### Network analysis of the identified clusters

We first constructed the complete network that consisted of all gene clusters from both transitions. Each node in this network represented a cluster of genes, and edges represented the genes that are shared between clusters. Since each gene is uniquely assigned to a single cluster in each transition, no two clusters from the same transition will ever be connected. Similarly, every gene is represented once, and only once, among all the edges.

The conserved network was constructed by looking at the Jaccard Index for each pair of clusters as a measure of similarity. We then calculated 1000 random cluster networks (each cluster had the same number of genes as the original), and calculated the Jaccard Indices of all node pairs. Our conserved network was then created by choosing only those edges that represent a Jaccard Index greater than 95% of all scores from the random networks.

## Acknowledgements

This work was supported by grant 1DP2GM105454-01 from the NIH, a Searle Scholar Award, a Klingenstein-Simons Fellowship Award in the Neurosciences, and a Pew Biomedical Scholar Award to DJCK. RL was funded by a Marie Curie international outgoing fellowship (PIOF-GA-2012-327992). PRO was supported by a Leon Levy Neuroscience Fellowship from the Leon Levy Foundation for Mind, Brain and Behavior. This is Clonal Raider Ant Project paper #10.

## Author contributions

PO and DK designed the study; RL and PO conducted the experiments and analyzed the data; RL wrote the manuscript with input from PO and DK; and DK supervised the study.

